# Efficient Zygotic Genome Editing via RAD51-Enhanced Interhomolog Repair

**DOI:** 10.1101/263699

**Authors:** Jonathan J. Wilde, Tomomi Aida, Martin Wienisch, Qiangge Zhang, Peimin Qi, Guoping Feng

## Abstract

Recent advances in genome editing have greatly improved knock-in (KI) efficiency^1–9^. Searching for factors to further improve KI efficiency for therapeutic use and generation of non-human primate (NHP) models, we found that the strand exchange protein RAD51 can significantly increase homozygous KI using CRISPR/Cas9 in mouse embryos through an interhomolog repair (IHR) mechanism. IHR is well-described in the context of meiosis^10^, but only occurs at low frequencies in mitotic cells^11,12^ and its existence in zygotes is controversial. Using a variety of approaches, we provide evidence for an endogenous IHR mechanism in zygotes that can be enhanced by RAD51. We show that this process can be harnessed for generating homozygous KI animals from wildtype zygotes based on exogenous donors and for converting heterozygous alleles into homozygous alleles without exogenous templates. Furthermore, we elucidate additional factors that contribute to zygotic IHR and identify a RAD51 mutant capable of insertion-deletion (indel)-free stimulation of IHR. Thus, our study provides conclusive evidence for the existence of zygotic IHR and demonstrates methods to enhance IHR for potential use in gene drives, gene therapy, and biotechnology.

---

We previously described a three-component CRISPR approach that employs Cas9 protein and chemically synthesized crRNA and tracrRNA for efficient KI of transgenes^1,2,9^. In addition, published work in rabbit embryos has shown significant enhancement of KI efficiency using RS-1, a chemical agonist of the homology-directed repair (HDR) pathway member RAD51^6^. We began by testing whether a combination of these techniques would promote high-efficiency knock-in of an autism-associated point mutation in *Chd2* (c.5051G>A; R1685H in human, R1684H in mouse; hereon referred to as *Chd2^R1684H^;* **Fig. 1a**) using a single-stranded oligonucleotide donor (ssODN). Previous studies have shown that KI efficiency is affected by the proximity of the Cas9 cut site and the insertion site^3^, so we chose a guide positioned where the cut site is directly adjacent to the desired G>A point mutation. Genotyping of mouse embryos after pronuclear injection and *in vitro* culture with or without RS-1 (7.5µM) showed a high KI efficiency at the targeted *Chd2* locus but revealed that RS-1 did not affect KI efficiency (**Extended Data Fig. 1**). Although the known mechanisms of RAD51 function suggest it may not promote knock-in with ssODNs, our previous success using exogenous Cas9 protein to promote high-efficiency KI^1,9^ and the fact that RS-1 effects are dosage-sensitive and uncharacterized in mouse embryos^6^ led us to hypothesize that the use of exogenous RAD51 protein in combination with three-component CRISPR could increase KI efficiency. We performed both pronuclear (PNI) and cytoplasmic (CPI) injections of crRNA, tracrRNA, Cas9, ssODN (30 ng/µL for PNI and 100 ng/µL for CPI), and RAD51 protein (10ng/µL) in mouse zygotes and genotyped the resulting F_0_ pups (example chromatograms in **Fig. 1b**). We observed overall KI efficiencies of 78% (7/9) and 56% (5/9) for CPI and PNI, respectively (**Fig. 1c**). Surprisingly, both CPI and PNI resulted in a homozygous KI efficiency of 56% (**Fig. 1d,** each 5/9). On-target editing was confirmed through the use of primers sitting outside of the ssODN homology arms and breeding of homozygous animals with wildtype C57BL/6N mice confirmed their homozygosity, as 36/36 F_1_ progeny from 6 mating pairs genotyped as *Chd2^R1684H/+^* (**Fig. 1e**).

**Figure 1.**
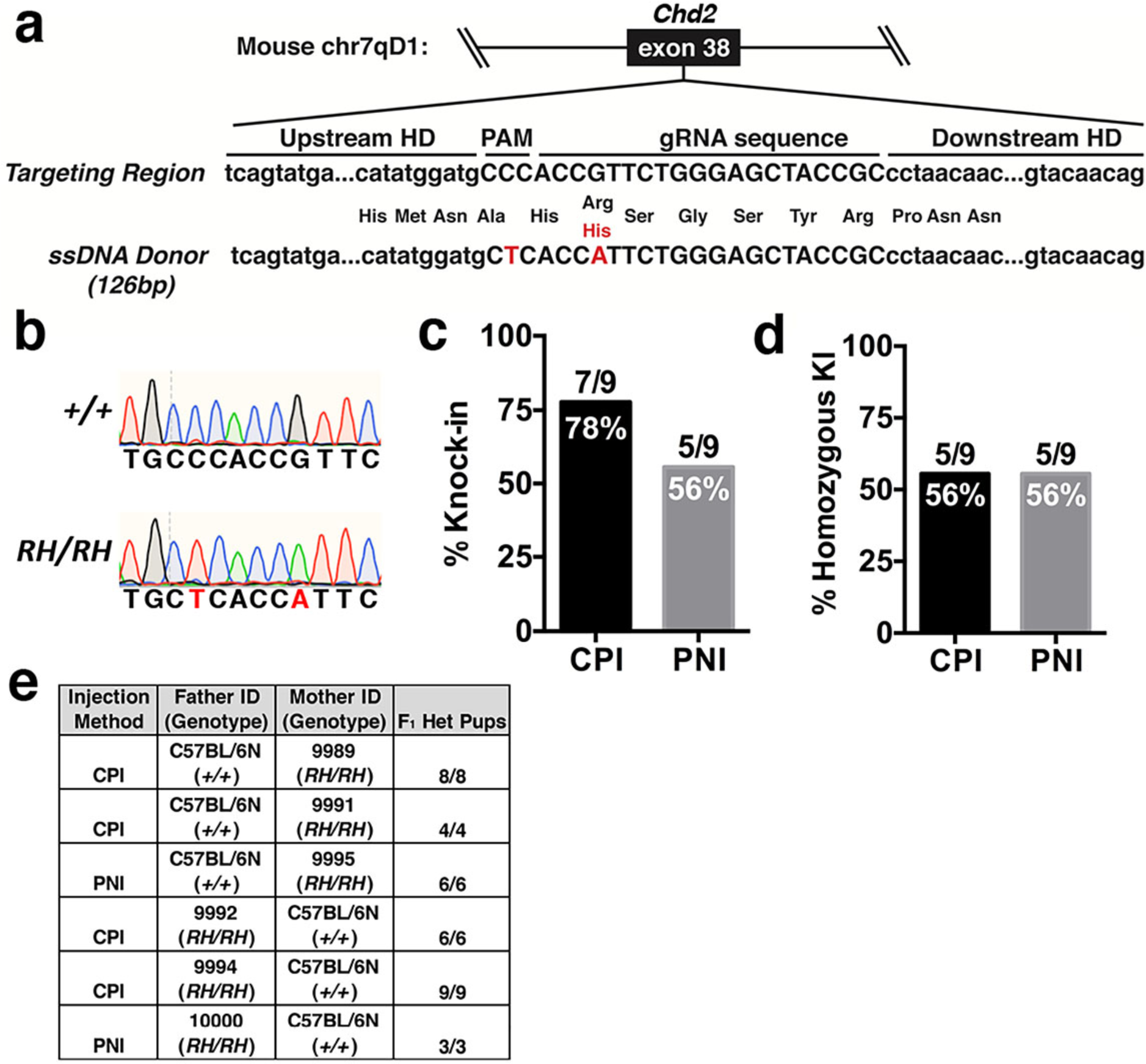
Efficient homozygous knock-in with RAD51. **a,** Schematic of the targeting locus and HR donor for generating *Chd2^R1684H^* mutant mice. HR donor contains both the c.5051G>A point mutation and a synonymous mutation that destroys the relevant PAM site (HD=homology domain). **b,** Example chromatograms of wildtype (top) and *Chd2^R1684H/R1684H^* animals (RH=R1684H). **c,** Overall KI efficiency observed in F_0_ pups derived from either cytoplasmic (CPI) or pronuclear (PNI) injection (pups with ≥1 KI allele/total pups). **d,** Homozygous KI rates observed in F_0_ pups generated by either CPI or PNI. **e,** Genotyping results from F_1_ pups derived from crosses between F_0_ *Chd2R1684H/R1684H* animals and wildtype C57BL/6N animals.

Since we did not initially generate *Chd2^R1684H^* mice from injections lacking RAD51, we performed additional injections to directly assess the ability of RAD51 to increase KI efficiency. We did not observe significant differences in efficiency between CPI and PNI (**Fig. 1c,d** and **Extended Data Table 1**), and therefore focused on our lab’s standard PNI injection strategy for the rest of this study. Zygotic injections into the paternal pronucleus were performed with and without RAD51, embryos were cultured for two days before lysis and DNA purification, and then nested PCR was performed. Sanger sequencing revealed that RAD51 increased overall KI efficiency (**Fig. 2a**, **left,** p=0.034, one-tailed chi-square test) and strongly increased homozygous KI efficiency (**Fig. 2a, middle**, includes mosaic embryos, p<0.0001, one-tailed chi-square test; **Fig. 2a, right**, excludes mosaic embryos, p=0.015). It is widely accepted that CRISPR/Cas9-mediated editing efficiency can be locus- and guide-dependent, so we tested the effects of RAD51 on KI efficiencies at a second genomic locus to confirm its effects. Using the same strategy employed for *Chd2*, we attempted to knock in an albinism-associated human mutation (c.265T>A; C89S; hereon referred to as *Tyr^C89S^*; schematic in **Fig. 2b**) in the *Tyr* gene^13^, which encodes for tyrosinase, the rate-limiting enzyme in melanin production. Albinism is a recessive disorder and, as such, only homozygous disruption of *Tyr* results in mice completely lacking pigment (**Fig. 2c**). Because homozygous indels in *Tyr* can also result in albinism, all F_0_ animals were genotyped to assess the effects of RAD51 on KI efficiencies (**Fig. 2d**) and phenotyped (**Extended Data Table 2)** to assess mosaicism. In the case of mosaic animals, genotyping was performed using tissue completely lacking melanin to assess the number of animals with a homozygous KI event (**Fig. 2e, middle)**. Injections without RAD51 showed a high overall KI efficiency and this efficiency was not significantly increased by the addition of RAD51 (**Fig. 2e, left**). However, our genotyping results showed that RAD51 enhances homozygous KI efficiency, with 44% of pups (including mosaics) having an albino phenotype resulting specifically from homozygous KI of the *Tyr^C89S^* allele with RAD51, as compared to only 12% without (**Fig. 2e, middle**, one-tailed chi-square test, p=0.0081). Furthermore, RAD51 co-injection increased the number of pure homozygous KI animals (**Fig. 2e, right,** one-tailed chi-square test, p=0.027), supporting our findings at the *Chd2* locus indicating that RAD51 can enhance homozygous KI.

**Figure 2.**
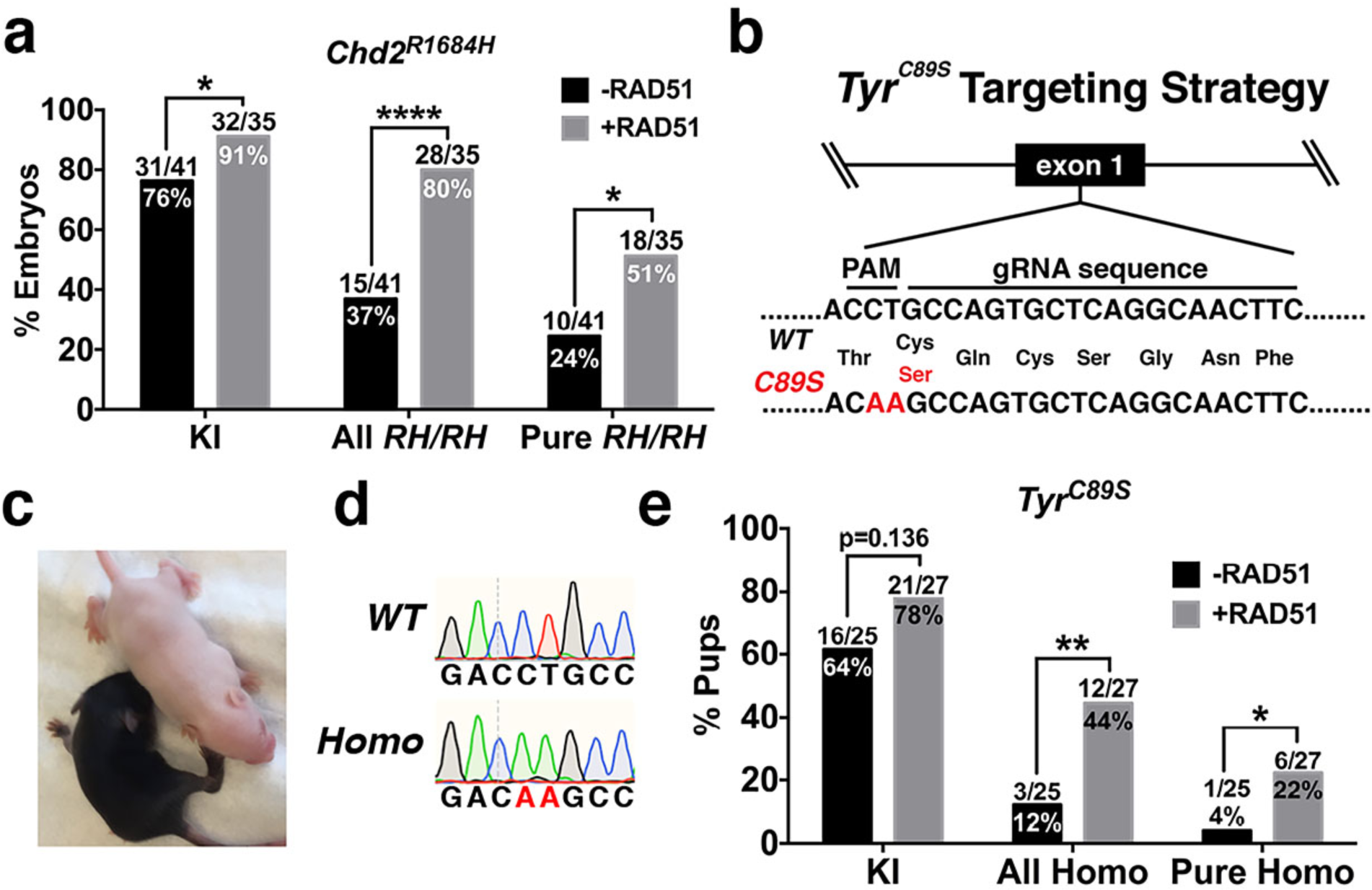
RAD51 enhances homozygous KI efficiency at multiple loci. **a,** Quantification of KI efficiency (left) and homozygous KI efficiency in embryos generated by *Chd2^R1684H^* PNI with or without RAD51. All *RH/RH* includes mosaic embryos and denotes embryos with >2:1 ratio of mutant allele to wildtype allele (KI: p=0.034, one-tailed chi-square test; All *RH/RH*: p<0.0001, one-tailed chi-square test; Pure *RH/RH*: p=0.015, one-tailed chi-square test). **b,** Schematic of knock-in strategy for the albinism-associated *Tyr^C89S^* mutation. **c,** Representative examples of *Tyr^+/+^* (black) and *Tyr^C89S/C89S^* (white) littermates derived from three-component CRISPR injections using exogenous RAD51 protein. **d,** Representative chromatograms from *Tyr^+/+^* (top) and *TyrC89S/C89S* animals. **e,** Genotyping of F_0_ animals generated by three-component CRISPR targeting the *Tyr^C89S^* locus with or without RAD51 (KI: p=0.136, one-tailed chi-square test; All Homo: p=0.0081, one-tailed chi-square test; Pure Homo: p=0.0272, one-tailed chi-square test).

Strikingly, RAD51 co-injection more than doubled the observed homozygosity at both the *Chd2* and *Tyr* loci, regardless of whether or not we took mosaicism into account (**Fig. 2a,e**). Previous attempts to increase HDR through pharmacological or genetic mechanisms have not shown such high homozygosity rates^3,7,14^, leading us to investigate the mechanism by which RAD51 can promote such an outcome. Our evidence that RAD51 only marginally increases the overall efficiency of ssODN-mediated knock-in (**Fig. 2a,e**) following PNI suggested that its effects on homozygosity do not come from increasing independent KI events on both alleles. Additionally, due to the random nature of indels produced by non-homologous end joining (NHEJ), we found it surprising that we observed multiple embryos from both *Chd2^R1684H^* and *Tyr^C89S^* injections whose genotyping results suggested the presence of the same indel on both ^alleles (**Extended Data Table 1,2**). Genotyping of F_1_ pups derived from a *Chd2*-targeted female^ with such a homozygous indel revealed that all pups were heterozygous for the same indel, confirming that the homozygosity of the founder (**Extended Data Fig. 2**). Although we could not completely rule out that these deletions were generated by MMEJ, which can result in stereotyped indels based on local homology^2,15^, the variety of observed indels and a lack of MMEJ sequence motifs at the R1684H editing site suggest that the homozygosity resulted from recombination between homologs that transferred the indel from one chromosome to the other.

More than 20 years ago, experiments in mouse ES cells found evidence of recombination events between homologous chromosomes^16,17^. While these interhomolog repair (IHR) events are well-understood in meiotic cells, their occurrence in zygotes and somatic cells is poorly described. Furthermore, experiments investigating IHR in somatic cells have shown that it occurs at very low frequencies^11,12^. However, several recent studies have described zygotic IHR events after induction of a targeted double-strand breaks (DSBs) by Cas9^18,19^. Based on these data, we hypothesized that RAD51 promotes homozygous KI through a similar mechanism. To directly test this, we attempted to generate homozygous *Chd2^R1684H^* embryos through editing of *Chd2^R1684/+^* embryos without the use of an exogenous donor. Using wildtype C57BL/6N eggs and sperm from homozygous *Chd2^R1684H^* male mice, we generated heterozygous zygotes via IVF and specifically targeted Cas9 to the wildtype maternal *Chd2* locus. As illustrated in **Figure 3a**, paternal PNI was performed on zygotes ~8 hours post fertilization (hpf) using a mixture of crRNA, tracrRNA, and Cas9 with or without RAD51. The *Chd2^R1684H^* allele contains mutations in both the guide sequence and its associated PAM and we confirmed through *in vitro* digestion assays that Cas9 is only capable of cutting the wildtype maternal allele (**Extended Data Fig. 3**). By omitting a donor template from the injection mixture, we were able to directly test both the rate of baseline CRISPR/Cas9-mediated IHR in zygotes and the ability of RAD51 to enhance such a mechanism. In agreement with previous work demonstrating IHR in human zygotes^18^, including mosaic embryos we observed IHR events in 26% of control-injected embryos (**Fig. 3b,c**). Strikingly, co-injection of RAD51 increased this rate to 74% (**Fig. 3c, left**, one-tailed chi-square test, p<0.0001). Analysis of the rates of embryos with pure, non-mosaic IHR events showed a similar increase in IHR rates in RAD51-injected embryos (**Fig. 3c, right**, one-tailed chi-square test, p=0.009), supporting the hypothesis that RAD51 can promote IHR in mouse zygotes.

**Figure 3.**
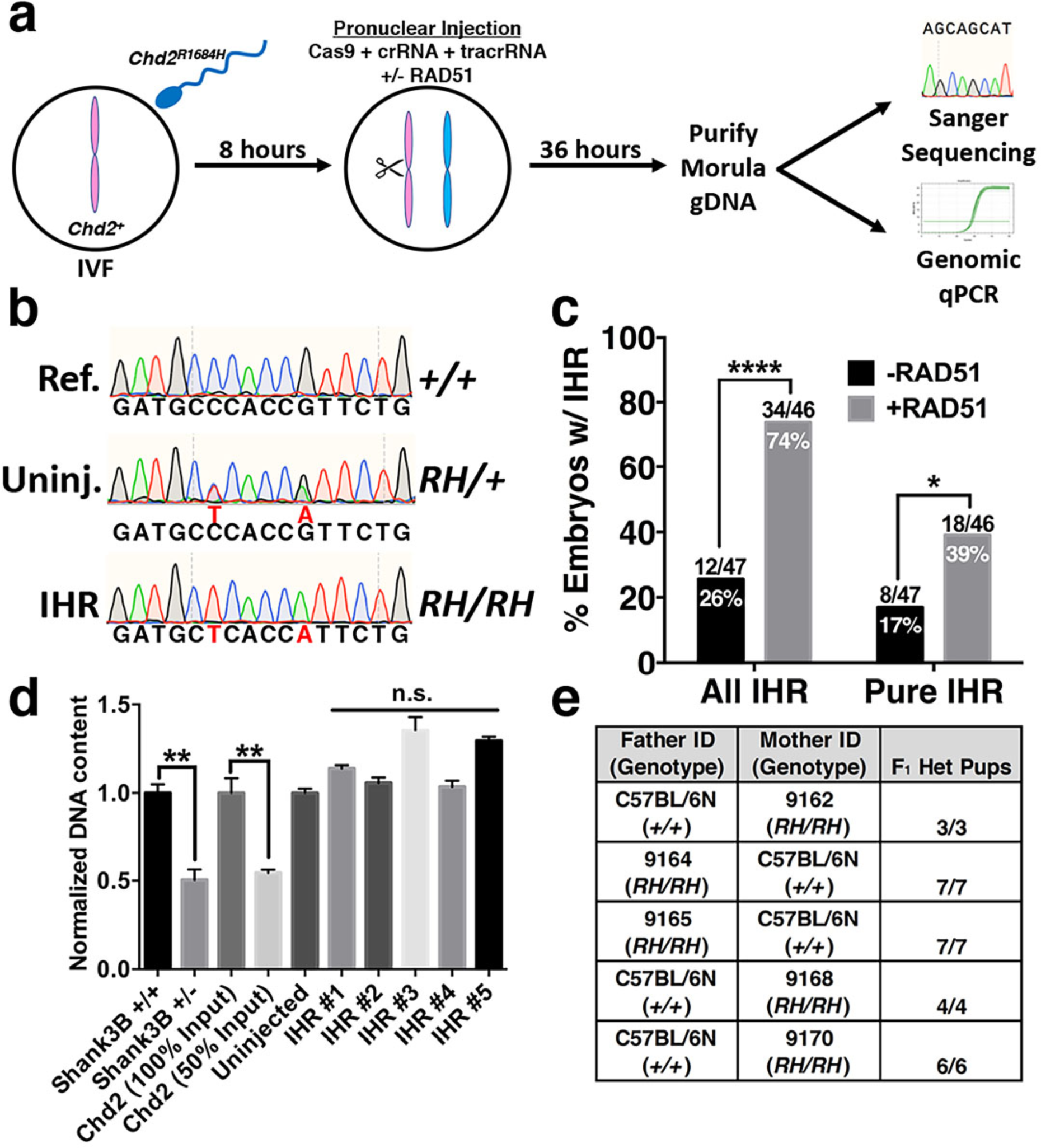
RAD51 enhances interhomolog repair. **a,** Schematic of strategy for testing RAD51-enhanced IHR. Wildtype C57BL/6N eggs were fertilized *in vitro* by sperm collected from male *Chd2^R1684H/R1684H^* mice and cultured for 8 hours. At 8hpf, PNI was performed with Cas9 protein, crRNA, and tracrRNA with or without RAD51 protein. Since *R1684H* mutants carry a mutation in the PAM associated with the crRNA utilized in these experiments, Cas9 is only capable of cutting the maternal allele. Injected embryos were then cultured for 48 hours and collected at morula stage. Half of the purified DNA was used for nested PCR and Sanger sequencing and the other half was used for multiplex PCR and subsequent qPCR to analyze genomic copy number at the *Chd2* editing locus. **b,** Representative chromatograms showing the wildtype reference sequence (Ref., top), an uninjected *Chd2^R1684H/+^* embryo generated by IVF (Uninj., middle), and a pure *Chd2^R1684H/R1684H^* homozygous mutant (IHR, bottom) generated via donor-free, RAD51-enhanced three-component CRISPR. **c,** Quantification of Sanger sequencing results from IVF-derived embryos co-injected with or without RAD51. ‘All IHR’ includes mosaic embryos and was determined by identifying embryos with >2:1 ratio of R1684H-to-wildtype alleles (one-tailed chi-square test, p<0.0001). Pure IHR refers to embryos without mosaicism (p=0.018, one-tailed chi-square test). **d,** Genomic qPCR targeting *Shank3B* using 180pg genomic DNA from wildtype and *Shank3B^+/-^* mice as a control for copy-number sensitivity (lanes 1-2, p=0.00295, unpaired t-test, t=6.47, df=4, *n*=3 technical replicates per sample, error bars=SEM). Lanes 3-4 illustrate genomic qPCR for the *Chd2^R1684H^* locus using 100% and 50% input of DNA derived from wildtype blastocysts for *Chd2* quantification (but 100% input for *Gapdh*) as a control for copy-number sensitivity at the locus (p=0.0099, unpaired t-test, t=5.355, df=3.302, *n*=4 technical replicates per condition, error bars=SEM). Lanes 5-10 illustrate genomic qPCR for the *Chd2^R1684H^* locus using DNA from an uninjected embryo and 5 randomly selected pure homozygous *Chd2^R1684H^* embryos (p>0.05, unpaired t-test, t=4.60, df=6, *n=4* technical replicates per sample, error bars=SEM)**. e,** F_1_ genotyping results of litters derived from crosses between wildtype C57BL/6N mice and 5 pure *Chd2^R1684H/R1684H^* F_0_ animals generated by IVF with RAD51.

Previous studies analyzing the properties of indels observed in single sgRNA-based CRISPR experiments have shown that indels large enough to disrupt our genotyping strategy are too infrequent to account for the loss-of-heterozygosity observed in our experiments^20^. Nonetheless, we performed multiple experiments to directly rule out the possibility of false positives resulting from monoallelic deletions on the maternal allele^21^. First, we developed and performed a multiplex genomic qPCR assay to analyze copy number at the targeted locus (**see Methods**). Using genomic DNA from wildtype mice and *Shank3B^+/-^* mice harboring a heterozygous deletion of exon 13 of *Shank3*^22^, we confirmed that our method was capable of identifying heterozygous deletions **(Fig. 3d)**. We then confirmed that PCR targeting the *Chd2^R1684H^* locus is sensitive enough to detect relevant changes in copy-number by halving the input DNA used for the assay (**Fig. 3d**). Finally, using this genomic qPCR strategy we were able to confirm normal copy-number at the R1684H locus in 5/5 randomly selected pure homozygous mutants (**Fig. 3d**). Next, we generated F_0_ *Chd2^R1684H/R1684H^* animals via the IVF strategy described in **Figure 3a** (including RAD51) and crossed homozygotes with wildtype C57BL/6N mice to generate F_1_ animals. Genotyping of F_1_ offspring confirmed that 100% were heterozygous for the R1684H mutation (**Fig. 3e**), supporting the conclusion that the F_0_ animals were homozygotes generated via IHR. Although we cannot rule out the possibility that a small fraction of embryos contain large hemizygous deletions, our results strongly support the conclusion that RAD51 is capable of stimulating highly-efficient zygotic IHR.

After fertilization, the maternal and paternal pronuclei remain physically separated until the completion of S-phase, at which time they fuse in preparation for mitosis. This poses a significant problem given that an IHR mechanism requires physical association of the maternal and paternal chromosomes and suggests that IHR must occur after S-phase. Because genotyping analyses of one-cell stage embryos are difficult to interpret due to failed PCR amplification of alleles harboring a Cas9-induced DSB, we chose to test the timing of zygotic IHR via immunocytochemistry (ICC) for RAD51 in uninjected, RAD51-injected, Cas9-injected (with *Chd2*-targeting crRNA), and Cas9- and RAD51-injected zygotes at three stages of the cell cycle: G1,, G2 (post S-phase), and M-phase (see **Methods**). Previous studies have shown that RAD51 staining is weak and/or diffuse in wildtype cells, but that strong RAD51 puncta appear in response to DSBs^23^. In agreement with our genotyping data and the hypothesis that IHR occurs after S-phase, we observed the appearance of RAD51 puncta solely during G2 (**Fig. 4a,b**). Although we did not observe RAD51 puncta in any uninjected G2 embryos, we were surprised to see several embryos injected solely with RAD51 that exhibited strong RAD51 foci (**Fig. 4b**). The appearance of such foci only after S-phase suggests that increasing levels of RAD51 can bias the cell toward RAD51-mediated repair over endogenous RAD52-dependent mechanisms for repairing replication-associated DSBs in mammals^24^. However, embryos injected with Cas9 showed a significantly increased frequency of RAD51 puncta and the highest proportion of positively-stained embryos was observed in the group injected with both Cas9 and RAD51 (**Fig. 4b, Extended Data Table 3**). Furthermore, embryos injected with RAD51 alone only exhibited single RAD51 puncta, but Cas-injected embryos were often observed with 2 or more puncta (**Extended Data Fig. 4**), indicative of IHR events on both maternal alleles, as well as at potential off-target Cas9-induced DSBs. While these findings suggest a model whereby RAD51 stimulates IHR in zygotes during G2, a deeper understanding of the pathways involved in IHR is still necessary.

**Figure 4.**
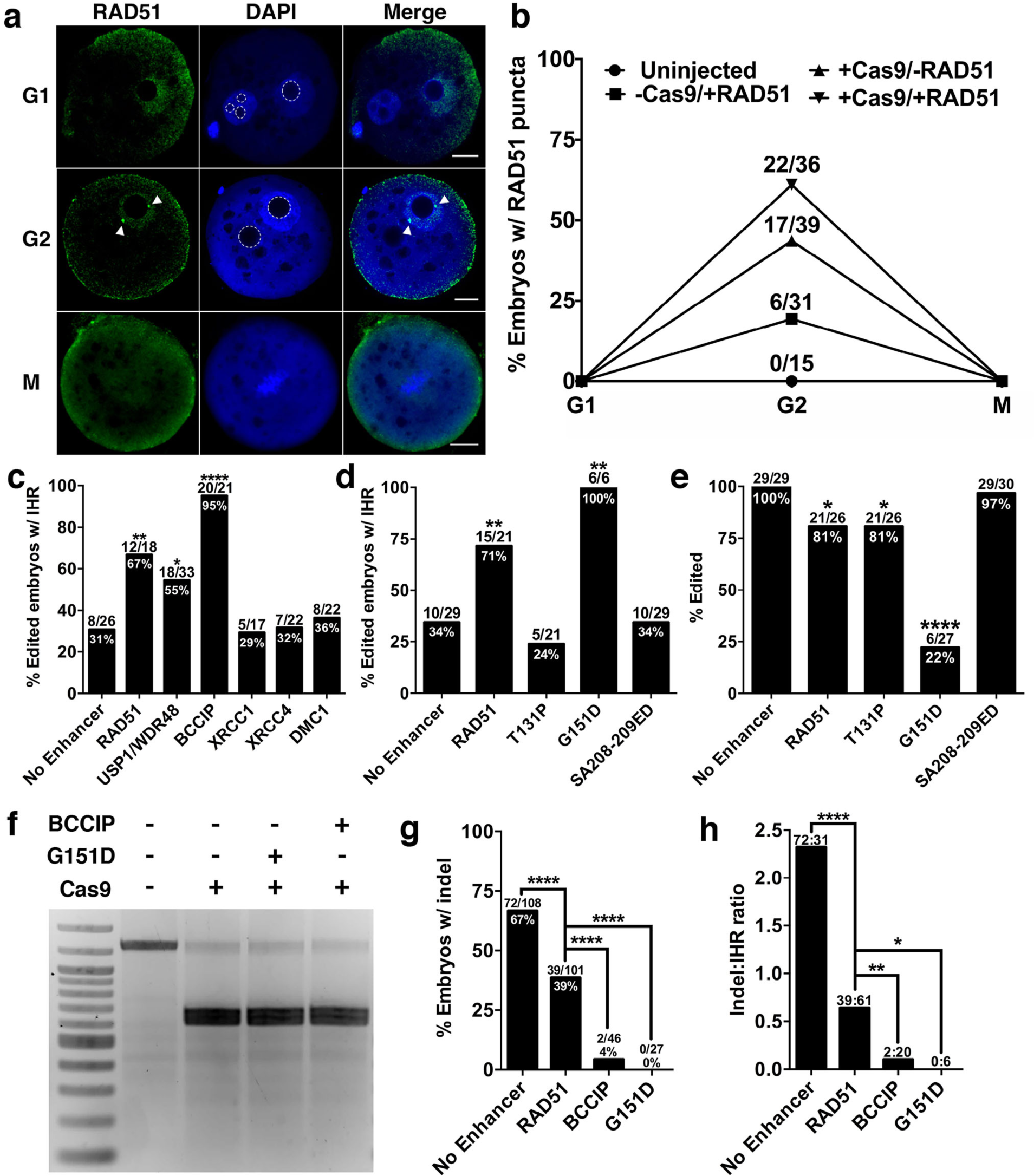
RAD51 G151D and BCCIP promote zygotic IHR and suppress indel formation. Representative images of RAD51 immunostaining from Cas9 + RAD51-injected embryos during G1, G2, and M-phase. Arrowheads indicate RAD51 puncta. Dotted lines highlight pronuclear nucleoli. **b,** Quantification of immunocytochemistry for RAD51 displaying the number of zygotes positive for RAD51 puncta during G1, G2, or M-phase. **c,** Quantification of IHR efficiency based on *Chd2^R1684H^* genotyping of blastocysts derived from embryos injected with Cas9/crRNA/tracrRNA and the indicated DSB repair-related proteins. Quantification includes mosaic embryos (one-tailed chi-square test). **d,** Quantification of IHR efficiency in blastocysts derived from Cas9/crRNA/tracrRNA-injected embryos co-injected with the indicated RAD51 variants (one-tailed chi-square test). **e,** Quantification of editing efficiency at the *Chd2^R1684H^* locus in embryos injected with Cas9/crRNA/tracrRNA and the indicated RAD51 variants (one-tailed chi-square test). **f,** *In vitro* Cas9 nuclease activity assay using a PCR amplimer of the *Chd2^R1684H^* locus and Cas9 alone or co-incubated with either RAD51 G151D or BCCIP. **g,** Quantification of the percent of total injected embryos carrying an indel after injection with the specified proteins. **h,** Quantification of the ratio of indel-positive embryos to IHR-positive embryos derived from injections with the specified proteins.

To gain a deeper understanding of the pathways involved in zygotic IHR, we performed a series of injections in Cas9-injected *Chd2^R1684H/+^* zygotes with multiple DSB repair- and homologous recombination (HR)-associated proteins: USP1/WDR48 (promotes HR via Fanconi-Anemia (FA) pathway^25^), BCCIP (promotes BRCA2-mediated HR^26-28^, participates in FA pathway^29^), XRCC1 (promotes MMEJ and co-localizes with RAD51^30,31^), XRCC4 (promotes NHEJ and regulator of V(D)J recombination^32,33^), and DMC1 (meiotic HR regulator^34^). XRCC1, XRCC4, and DMC1 were not capable of inducing IHR at rates above baseline (**Fig. 4c**), confirming the specificity of RAD51 and indicating that zygotic IHR pathways do not utilize proteins typically related to HR-independent DSB repair. Co-injection of USP1/WDR48, on the other hand, was capable of significant enhancement of IHR (**Fig. 4c**). Strikingly, BCCIP exhibited a significantly stronger effect on IHR rates than USP1/WDR48 (one-tailed chi-square test, p=0.0014), with 95% of edited embryos showing IHR (**Fig. 4c**). Since BCCIP-mediated activation of BRCA2 promotes RAD51 activation, these data suggest that proteins capable of directly affecting RAD51 activity are likely to promote high-efficiency IHR. Additionally, the ability of the FA pathway members BCCIP, USP1/WDR48, and RAD51 to enhance zygotic IHR implicates the FA pathway in this process.

*RAD51* mutations have been identified in multiple human cancers and Fanconi-Anemia and several of these mutations have been studied in-depth. In an effort to better understand the role of RAD51 in IHR, we co-injected three different recombinant RAD51 mutant proteins with Cas9 and *Chd2^R1684H^* crRNA into IVF-generated *Chd2^R1684H/+^* embryos. The mutants were as follows: T131P (deficient in interstrand crosslink repair (ICL), but not HR^35,36^), SA208-209ED (disrupted BRCA2-association^37^), and G151D (gain-of-function mutant with increased strand exchange activity^38^). T131P injection was not capable of increasing IHR frequency compared to Cas9-only embryos (**Fig. 4d**), indicating that RAD51 function associated with ICL repair is required for the induction of IHR. Likewise, SA208-209ED showed unchanged IHR rates compared to Cas9-only embryos (**Fig. 4d**), supporting a role for BRCA2 in RAD51-mediated promotion of IHR. Surprisingly, in 27 zygotes injected with the RAD51 gain-of-function variant, we did not record a single indel event and observed IHR in all edited cells (**Fig. 4d**). However, the apparent editing efficiency dropped to 22% (**Fig. 4e**), suggesting that G151D either directly inhibits Cas9 nuclease activity or stimulates IHR at such high levels that the replicated wildtype maternal allele is often used as a template. To test whether G151D can directly inhibit Cas9 activity, we performed *in vitro* Cas9 digestion assays with and without co-incubation with G151D and did not observe any direct inhibition of Cas9 activity (**Fig. 4f**). Interestingly, we observed a similar decrease in editing efficiency with BCCIP-injected embryos that was not due to direct inhibition of Cas9 activity (**Fig. 4f, Extended Data Fig. 5**). Additional analyses of overall indel rates and the indel-to-IHR ratios in embryos injected with RAD51, BCCIP, or RAD51 G151D revealed a drastic decrease in both rates compared to control embryos (**Fig. 4g,h**). Furthermore, both BCCIP and RAD51 G151D embryos showed significant decreases in these rates compared to embryos injected with wildtype RAD51. Our previous experiments demonstrated that IHR occurs after S-phase in zygotes, when a second wildtype allele is present to serve as an IHR template. Therefore, our data argue that rather than inhibiting Cas9-mediated DSBs, BCCIP and the RAD51 G151D mutant stimulate IHR with nearly 100% efficiency and indel-to-IHR ratios near zero.

Our data demonstrate that RAD51, the BRCA2-interacting protein BCCIP, and the FA pathway-associated protein complex of USP1 and WDR48 are capable of significantly enhancing the rate of zygotic interhomolog repair. Our mechanistic characterization of IHR in zygotes suggests a model whereby Cas9-induced DSBs are repaired through the use of a homologous chromosome as a template during G2. At this time, the maternal and paternal genomes have already undergone replication and the pronuclei undergo fusion prior to the start of M-phase. This creates a significant hurdle for pure, non-mosaic IHR, as the cell must now edit two alleles instead of one. This is highlighted by the high degree of mosaicism observed in embryos exhibiting IHR (**Extended Data Fig. 6**). Interestingly, although BCCIP-injected embryos exhibited a significant increase in mosaicism, only 1 out of 14 BCCIP-injected embryos with mosaic IHR was mosaic with an indel (**Extended Data Fig. 6c,d**), suggesting that BCCIP protects against indels by greatly increasing the likelihood of repairing DSBs through IHR instead of NHEJ. Together, our mosaicism analyses support our ICC results showing that zygotic IHR occurs following pronuclear fusion when there are multiple IHR templates available for repair.

Single-cell RNA-sequencing has shown that *Rad51* mRNA is present in zygotes at relatively low levels (FPKM <50)^39^. Nonetheless, the existence of endogenous RAD51 in zygotes begs the question of exactly how exogenous RAD51 enhances IHR. We hypothesize based on the data presented in this paper that increased concentrations of RAD51 in zygotic pronuclei increase the probability of RAD51 locating and binding to a DSB. While previous work has shown that accumulation of endogenous RAD51 at zygotic DSBs can occur in genetic models of genomic instability, we did not observe any RAD51 puncta in uninjected cells at any stage of the cell cycle. However, injection of RAD51 protein alone revealed a small number of cells with RAD51 puncta (**Fig. 4b**), suggesting that increasing RAD51 levels is capable of biasing repair of endogenous DSBs toward RAD51-dependent pathways. The highly-efficient, indel-free editing observed in cells injected with the G151D mutant further supports this mechanism where increased concentrations of RAD51 promote IHR by increasing the likelihood of RAD51 binding to a DSB. Although we injected the same concentration of G151D into cells as we did wildtype RAD51, the G151D mutant has a faster on-rate for binding DSBs and binds DSBs more stably due to its increased strand-exchange activity^38^ and decreased rate of ATP hydrolysis^40^, respectively. This could help explain why we do not observe indels in these cells, as G151D is likely capable of binding Cas9-induced DSBs faster and for longer periods of time than other DNA repair molecules after the completion of S-phase.

Our insights into the basic mechanisms of IHR in zygotes suggest that enhanced IHR has the potential to overcome current problems limiting the feasibility of gene drive strategies. Gene drives are designed to edit wild populations for disease prevention and agricultural purposes by accelerating the spread of alleles conferring desired traits. They consist of a single locus encoding a gene of interest, Cas9, and an sgRNA targeting the same wildtype locus. In heterozygotes, expression of Cas9 and the sgRNA targets Cas9 to the wildtype allele, inducing a DSB that can be repaired through IHR to generate cells homozygous for the gene drive cassette. This conversion of heterozygotes to homozygotes is key to the success of gene drives, but recent work has demonstrated the rapid development of gene drive resistance through the production of undesired indels that prevent cutting and subsequent IHR^41,42^. Therefore, our discovery that BCCIP and RAD51 G151D promote nearly indel-free IHR in zygotes has enormous potential for overcoming the barriers to successful gene drive applications. Furthermore, the same advantages of indel-free IHR could be used to develop safe gene therapies for autosomal dominant disorders caused by point mutations and small indels. Given that we still know so little about endogenous mechanisms of IHR, the potential impact of our findings suggests that additional studies of IHR in zygotes and somatic cells will be fundamental to the development of improved tools and strategies for gene editing.

## Methods

### Data Availability

The raw data supporting the findings in this manuscript can be obtained from the corresponding author upon reasonable request.

### Animal Care and Use

All mouse work was performed with the supervision of the Massachusetts Institute for Technology Division of Comparative Medicine (DCM) under protocol 0416-024-19, which was approved by the Committee for Animal Care (CAC). All procedures were in accordance with the guidelines set forth by the Guide for Care and Use of Laboratory Animals, National Research Council, 1996 (institutional animal assurance no. A-3125-01). All embryos injected for the experiments described in this manuscript were on a C57BL/6NTac background (Taconic; referred to as C57BL/6N in paper).

### Preparation of Injection Mixtures

tracrRNA, crRNAs, and ssODNs were synthesized by Integrated DNA Technologies (see **Supplementary Table 1** for sequences). All injection mixtures were prepared in a final volume of 50µL according using the same protocol. Using RNase-free water, reagents, and consumables, crRNA (final concentration 0.61µM), tracrRNA (final concentration 0.61µM), and ultrapure Tris-HCl, pH7.39 (final concentration 10mM, ThermoFisher) were mixed and incubated at 95°C for 5 minutes. The mixtures were cooled to room temperature for 10 minutes on the benchtop and then EnGen Cas9 NLS, *S. pyogenes* (New England Biolabs) was added to a final concentration of 30ng/µL. The mixtures were incubated at 37°C for 15 minutes before adding any remaining components: ssODN (final concentration 30ng/µL), RAD51 (Creative Biomart, final concentration 10ng/µL). Injection mixtures were stored on ice and briefly heated to 37°C prior to injection. For experiments utilizing RS-1, embryos were cultured in KSOM-AA (EMD Millipore MR-121-D) with 7.5µM RS-1 (Sigma) dissolved in DMSO or DMSO (Sigma, 1:1000) for 24 hours, washed, and cultured in EmbryoMax FHM HEPES Buffered Medium (Sigma) until collection for genotyping.

### Recombinant Proteins

For experiments using wildtype recombinant proteins, the following proteins were used: RAD51 (Creative Biomart RAD51-134H), USP1/WDR48 (Creative Biomart USP1&WDR48-1067H), BCCIP (Origene TP303061), XRCC1 (TP304952), XRCC4 (Origene TP312684), DMC1 (Origene TP318311). For experiments using mutant RAD51, custom RAD51 preparations were produced by Creative Biomart according to their standard protocol for producing the wildtype RAD51 used in all other experiments. DNA sequences used for cloning the mutant forms of *Rad51* are described in **Supplementary Table 2**. All RAD51 mutants were modeled in human *RAD51* transcript variant 4 (NCBI accession NM_001164269), as this is the transcript variant of the wildtype RAD51-134H protein used in all other experiments.

### Natural Mating for Zygotic Injections

Female mice (4-5 weeks old, C57BL/6N) were superovulated by IP injection of PMS (5 IU/mouse, three days prior to microinjection) and hCG (5 IU/mouse, 47 hours after PMS injection) and then paired with males. Plugged females were sacrificed by cervical dislocation at day 0.5pcd and zygotes were collected into 0.1% hyaluronidase/FHM (Sigma). Zygotes were washed in drops of FHM and cumulus cells were removed. Zygotes were cultured in KSOM-AA for one hour and then used for microinjection.

### In Vitro Fertilization for Zygotic Injections

*In vitro* fertilization was performed using FERTIUP^®^ Mouse Preincubation Medium and CARD MEDIA (Kyudo Company) according to the manufacturer’s protocol. Non-virgin *Chd2^R1684H/R1684H^* males that were >8 weeks old were used as sperm donors. Following IVF, embryos were cultured for 8 hours and then injected using the PNI protocol described below.

### Zygotic Microinjections

For all experiments except the CPI experiments described in **Fig. 1**, the male pronucleus was injected. Injections performed for the CPI portion of **Fig. 1** were targeted to the cytoplasm. All microinjections were performed using a Narishige Micromanipulator, Nikon Eclipse TE2000-S microscope, and Eppendorf 5242 microinjector. Individual zygotes were injected with 1-2pL of injection mixture using an “automatic” injection mode set according to needle size and adjusted for clear increase in pronuclear volume. Following injections, cells were cultured in KSOM-AA until collection for genotyping. For experiments giving rise to F_0_ animals, embryos were surgically implanted into pseudopregnant CD-1 females (Charles River Laboratories, Strain Code 022) 24-hours post-injection and allowed to develop normally until natural birth.

### Embryo Collection and DNA Purification

Embryos were collected at morula-to-blastocyst stage in 4µL nuclease-free water. After collection, 4µL of 2X embryo digestion buffer was added to each sample (final concentrations: 125µg/mL proteinase K, 100mM Tris-HCl pH 8.0, 100mM KCl, 0.02% gelatin, 0.45% Tween-20, 60µg/mL yeast tRNA) and embryos were lysed for 1 hour at 56°C. Proteinase K was inactivated via incubation at 95°C for 10 minutes and DNA was stored at −20°C until use.

### Tail DNA Purification

Tail tissue (~0.5cm length) was collected from individual animals, placed in 75µL alkaline lysis buffer (25mM NaOH, 0.2mM EDTA), and incubated at 95°C for 30 minutes. Digestion was stopped via addition of 75µL neutralization buffer (40mM Tris-HCl pH 5.0). Samples were stored at 4°C until genotyping.

### Chd2^R1684H^ PCR

*Chd2^R1684H^*genotyping was performed via a two-step, nested approach. Except for experiments described in **Figure 3**, all 8µL of embryo DNA were used as input for the initial, long round of PCR. Genotyping of mouse pups and adult animals was performed using 2µL of purified tail DNA. For all embryo genotyping, reactions mixtures were set up in a UV-sterilized laminar flow hood using sterile reagents and filter tips to avoid contamination. All PCR was performed using Biolase DNA Polymerase (Bioline), fresh aliquots of dNTPs (New England Biolabs), and 2% DMSO (final concentration). The first PCR reaction of the nested PCR was performed using the C2RH-Long_F/R primer pairs listed in **Supplementary Table 3**. Primer pairs C2RH-Long_F1/R1 and C2RH-short_F1/R1 were used for **Figures 1–3**, as well as

**Extended Data Figures 1 and 2**. All other experiments utilized primer pairs C2RH-Long_F2/R2 and C2RH-short_F2/R2, which was designed to amplify a larger genomic region as a way to further rule out large deletions that could disrupt PCR primer binding sites. 20 cycles of amplification were performed using 30 second extension and annealing times and an annealing temperature of 67°C. Nested PCR was then performed using 2µL of the initial 25µL PCR reaction as input. The C2RH-Short_F/R primer pairs listed in **Supplementary Table 3** were used for this PCR, with sets 1 and 2 being used as described above. Thirty-five cycles of amplification were performed using 30 second extension and annealing times and an annealing temperature of 68°C.

### Tyr^C89S^ PCR

Initial PCR was performed using 2µL purified tail DNA. The longer amplification of the nested PCR reaction was performed using the Tyr-Long_F/R primer pair listed in **Supplementary Table 3**. PCR was performed using Biolase DNA polymerase (Bioline) and a final concentration of 2% DMSO. Twenty cycles of PCR were run with 30 second annealing and extension steps and an annealing temperature of 66°C. Nested PCR was performed using 2µL of the initial 25µL PCR reaction and the Tyr-Short_F/R primer pair listed in **Supplementary Table 3**. The same PCR conditions were used for the nested PCR as were used for the initial PCR.

### Sanger Sequencing

For all sequencing reactions, 5µL of PCR product was mixed with 3µL dH_2_O and 2µL ExoSAP (Exonuclease I, NEB #M0293 and rSAP, NEB #M0371). Products were incubated at 37°C for 30 minutes to degrade primers and dephosphorylate dNTPs and enzymes were then heat inactivated at 80°C for 15 minutes. 2.5µL dH_2_O and 2.5µL 10µM sequencing primer were then added to each mixture and samples were submitted to Genewiz for Sanger sequencing. Sequence files (.ab1) were blinded using a freely-available Perl script (https://github.com/jimsalterjrs/blindanalysis) and analyzed using SnapGene. To identify IHR events in mosaic samples, we monitored the genotype at the R1684H locus, as well as the PAM site and compared the major and minor peak heights using 4Peaks (Nucleobytes). Sequences showing >2:1 ratio of major allele to minor allele at both sites were called IHR-positive, but mosaic. Sequencing primers used were: Chd2-Short_R1, Chd2-Short_F2, and Tyr-Short_F (see **Supplementary Table 3**).

### Total Protein Staining and Western Blotting

1µg of each recombinant protein was mixed with 2X Laemmli Sample Buffer (Bio-Rad) and heated to 95°C for 5 minutes. After brief centrifugation, samples were loaded into 4-15% Mini-PROTEAN TGX Pre-Cast gels (Bio-Rad) along with 5µL PrecisionPlus Protein Kaleidoscope Prestained Protein Standards (Bio-Rad) and run at 200V for 40 minutes in 1X Tris/Glycine/SDS running buffer. Protein was transferred to nitrocellulose at 100V for 1 hour at 4°C and total protein was visualized using REVERT Total Protein Stain (LI-COR Biosciences) according to the manufacturer’s protocol. For RAD51 Western blotting in **Supplementary Figure 1a**, REVERT Total Protein Stain was reversed according to the manufacturer’s protocol, the membrane was blocked at room temperature for 1 hour in TBST containing 5% non-fat dry milk. Primary antibody (Rb anti-RAD51, Abcam ab63801, 1:500) was diluted in blocking buffer and applied to the membrane overnight at 4°C. Following washes with TBST, secondary antibody (IRDye 800CW Gt anti-rabbit IgG H+L, LI-COR 925-32211, 1:15,000) was diluted in blocking buffer and applied to the membrane for 1 hour at room temperature in the dark. The membrane was then washed and imaged. For all experiments, imaging was performed on an Odyssey CLx Imaging System with ImageStudio (LI-COR Biosciences). Coomassie staining and Western blots in **Supplementary Figure 1b** were performed by Creative Biomart.

### In vitro Cas9 Digestion Assays

Wildtype or *Chd2^R1684H/R1684H^* genomic DNA was used as input for PCR using the conditions described above and the primers specified in **Supplementary Table 3**. After confirmation of a single band via gel electrophoresis, the PCR reactions were purified using the DNA Clean & Concentrator kit (Zymo) according to the manufacturer’s instructions. crRNA and tracrRNA were diluted 1µM in Nuclease-free Duplex Buffer (IDT) and incubated at 95°C for 5 minutes before being allowed to cool to room temperature on the bench. Digestion mixtures were made in separate tubes by combining nuclease-free water, 10X Cas9 Nuclease Reaction Buffer (final concentration 1X, New England Biolabs), 1µL of the cooled crRNA/tracrRNA duplex, and, if necessary, 1µL of EnGen Cas9 NLS, *S. pyogenes* (New England Biolabs). These mixtures were heated to 37°C for 10 minutes before adding 250ng PCR product and, if necessary, RAD51 G151D (10ng/µL final concentration, custom-produced by Creative Biomart) or BCCIP (10ng/µL final concentration, Origene TP303061). Digestion was performed at 37°C for 1 hour, Cas9 was denatured at 95°C, 6X Purple Gel Loading Dye (New England Biolabs) was added to a final concentration of 1X, and the samples were allowed to passively cool to room temperature. After cooling, reactions were separated via electrophoresis with 2% agarose gel, post-stained with GelRed Nucleic Acid Gel Stain (Biotium) according to manufacturer’s instructions and visualized using an InGenius Gel Documentation System (Syngene).

### Genomic qPCR

To determine copy number at specific genomic loci, we developed a strategy utilizing multiplex nested qPCR. We first performed multiplex amplification of both the edited *Chd2* region and a region of the *Gapdh* promoter, which is located on a different chromosome, in a short round of PCR (10 cycles). In the case of our *Shank3B* control experiment, we attempted to match the input DNA concentration used for experiments utilizing DNA from cultured embryos. Based on an estimation of ~6pg DNA per cell and a PCR input of ~30 cells, we used 180pg *Shank3B^+/+^* or *Shank3B^+/-^* genomic DNA per initial reaction using the Shank3-Long-F/R and Gapdh-Long_F/R primer pairs described in **Supplementary Table 3**. We then used the PCR products from the initial multiplex PCR as input for nested qPCR using the Shank3-Short_F/R and Gapdh-Short_F/R primer pairs with Sso Advanced SYBR Green Supermix (Bio-Rad). qPCR was performed on a CFX96 Touch Real-Time PCR Detection System (Bio-Rad) and analyzed using CFX Maestro software (Bio-Rad). *Shank3B* signal was normalized to *Gapdh* signal to control for input and a second diploid locus. In a second control experiment to test qPCR sensitivity at the *Chd2* locus, we isolated DNA from wildtype blastocysts and performed multiplex PCR for *Chd2* using the C2RH-Long_F1/R1 and Gapdh-Long_F/R primer pairs listed in **Supplementary Table 3**. We then performed genomic qPCR using the same input amount for *Gapdh* nested qPCR (Gapdh-Short_F/R primers), but either 1X or 0.5X input for the *Chd2* nested qPCR (Chd2-Short_F1/R1 primers). Finally, we used the same strategy to perform nested genomic qPCR using genomic DNA isolated from 5 randomly-selected blastocysts that had genotyped positive for interhomolog repair with no evidence of mosaicism.

### Immunocytochemistry

Zygotes were collected during G1 (9hpf, 1hpi, PN2), G2 (14.5hpf, 6.5hpi, PN5), or M-phase (16hpf, 8hpi) and fixed overnight at 4°C in 4% PFA + 0.1% Tween-20 in PBS and then briefly transferred to acidified Tyrode’s (Sigma T1788) to remove the zona. Collection times and cell cycle stages were determined empirically based on pronuclear/nuclear morphology^43^. After washing in PBS + 0.1% Tween-20 (PBST), zygotes were permeabilized in PBS + 1% Triton X-100 for 1 hour at 4°C and then blocked for 1 hour at room temperature in blocking solution (PBS + 3% BSA + 5% normal goat serum). Primary antibody (rabbit anti-RAD51, Abcam ab63801, 1:500) was diluted in blocking solution and applied to coverslips overnight at 4°C. Cells were washed 3 times in PBST before application of secondary antibody (goat anti-rabbit IgG conjugate, Alexa Fluor 488, ThermoFisher A-11008) diluted in blocking solution for 1 hour at room temperature. Nuclei were counterstained with DAPI and coverslips were mounted to slides with Fluoromount Aqueous Mounting Medium (Sigma F4680). Cells were imaged on an Olympus Fluoview FV1000 confocal microscope with a 60X oil immersion objective and variable digital zoom. For all cells, we acquired z-stacks incorporating both pronuclei. Raw image files were exported to FIJI^44^ for Z-projection (sum slices), channel splitting, and scale bar generation.

### Statistical Analyses

Statistical analyses were performed using Prism 6 (Graphpad). For chi-square analyses, one-tailed tests were used in cases where we were testing whether a factor could specifically increase interhomolog repair. For all other experiments, two-tailed analyses were performed. Details on statistical tests performed for each experiment are provided in text and figure legends. All graphs display mean±SEM. *p<0.05, **p<0.01, ***p<0.001, ****p<0.0001

## Supplementary Information

Additional data and sequences of oligos, proteins, and primers used in this study can be found in Supplementary Information.

## Acknowledgements

The experiments described in this manuscript were funded by the Tan-Yang Center for Autism Research at MIT and the Poitras Center for Affective Disorders Research at MIT. We would like to thank Tetsushi Sakuma and Takashi Yamamoto of Hiroshima University for sharing their knock-in enhancer screening data. We would also like to thank Ricardo del Rosario for assistance with analysis of published single-cell RNA-seq data. Finally, we are grateful for the helpful discussion and insight provided by Charles Jennings, Sabbi Lall, and members of the Feng lab.

## Author Contributions

J.J.W., T.A., M.W., Q.Z., and G.F. designed experiments. J.J.W. and P.Q. performed experiments. J.J.W., T.A., M.W., and G.F. analyzed data. J.J.W., T.A., and G.F. prepared the manuscript.

## Reprints

Reprints and permission information is available at www.nature.com/reprints.

## Materials and Correspondence

Correspondence and requests for materials should be addressed to G.F. at

---

**Extended Data Figure 1.**
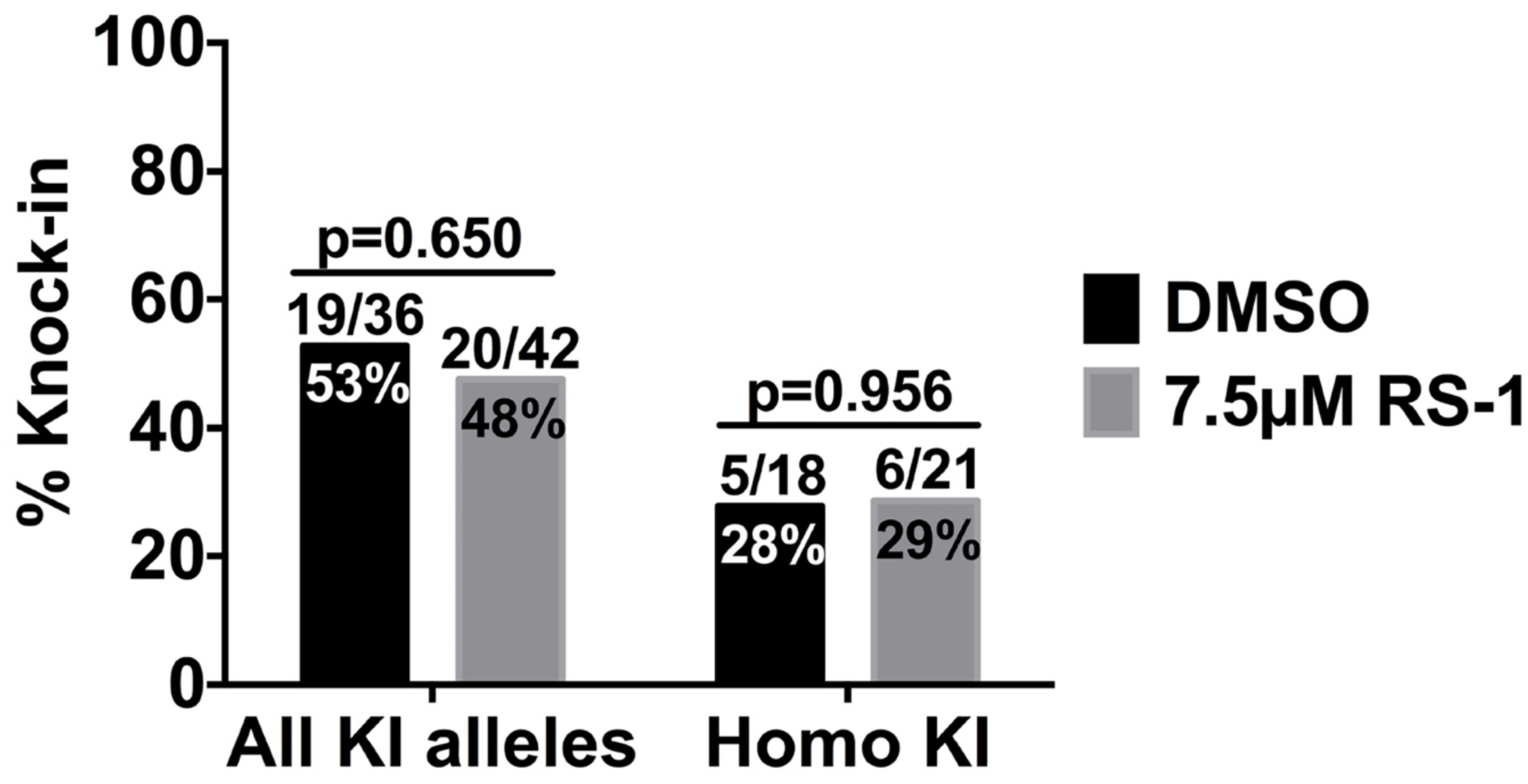
RS-1 does not alter KI efficiency with ssODNs in mouse zygotes. *Chd2^R1684H^* KI and homozygous KI efficiency in embryos cultured in either DMSO or 7.5µM RS-1 for 24 hours (two-tailed chi-square test).

**Extended Data Figure 2.**
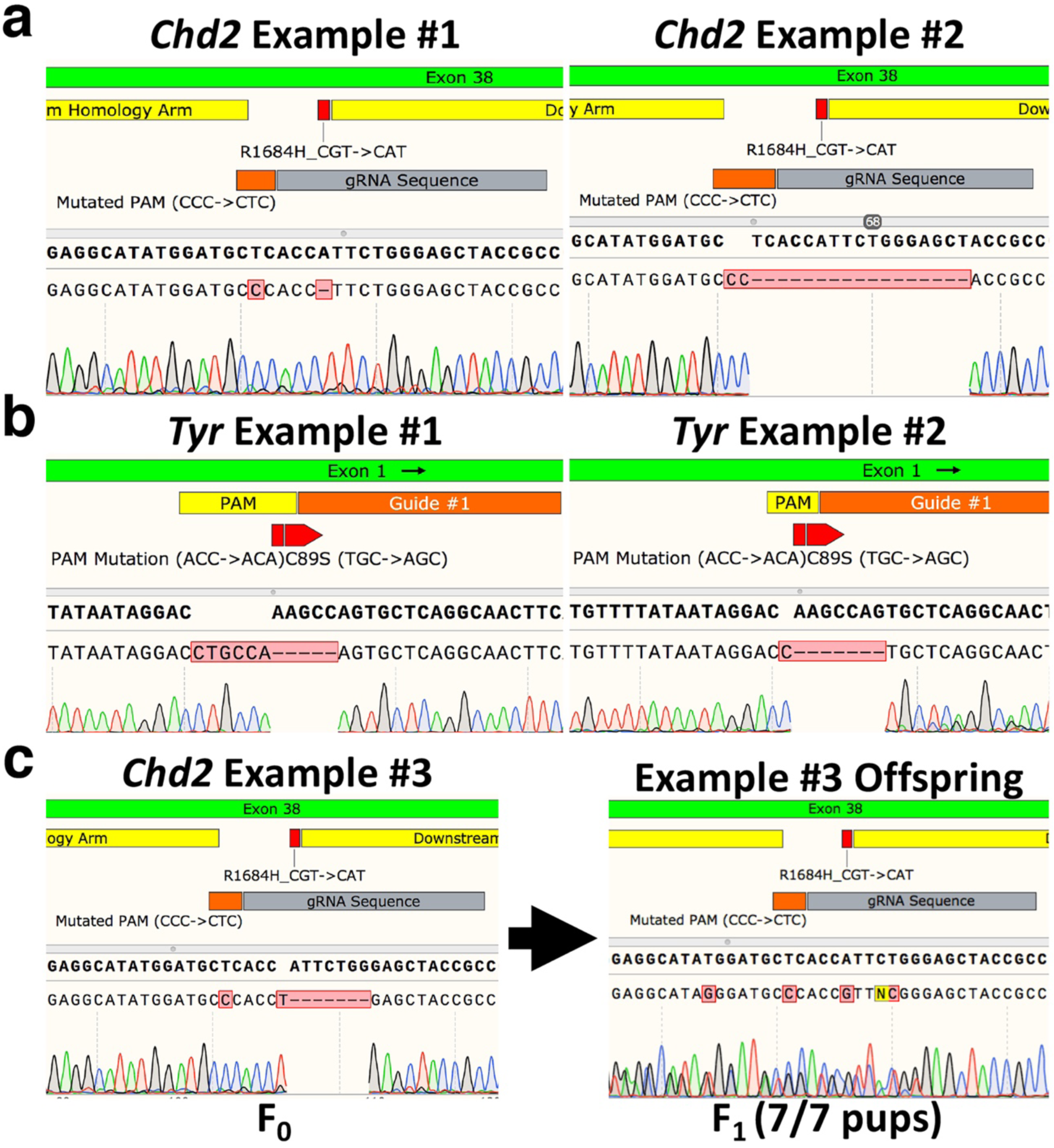
RAD51 stimulates the generation of homozygous indels. **a,** Representative Sanger sequencing traces of putative homozygous indels observed in F_0_ animals obtained from *Chd2^R1684H^* knock-in experiments. **b,** Representative Sanger sequencing traces of putative homozygous indels observed in F_0_ animals obtained from *Tyr^C89S^* knock-in experiments. **c,** Genotyping traces for a putative homozygous *Chd2* indel and an F_1_ offspring from a cross between the founder and a wildtype C57BL/6N mate. The F_1_ genotyping, which was consistent for all 7 pups in the litter, indicates heterozygous presence of the indel observed in the founder.

**Extended Data Figure 3.**
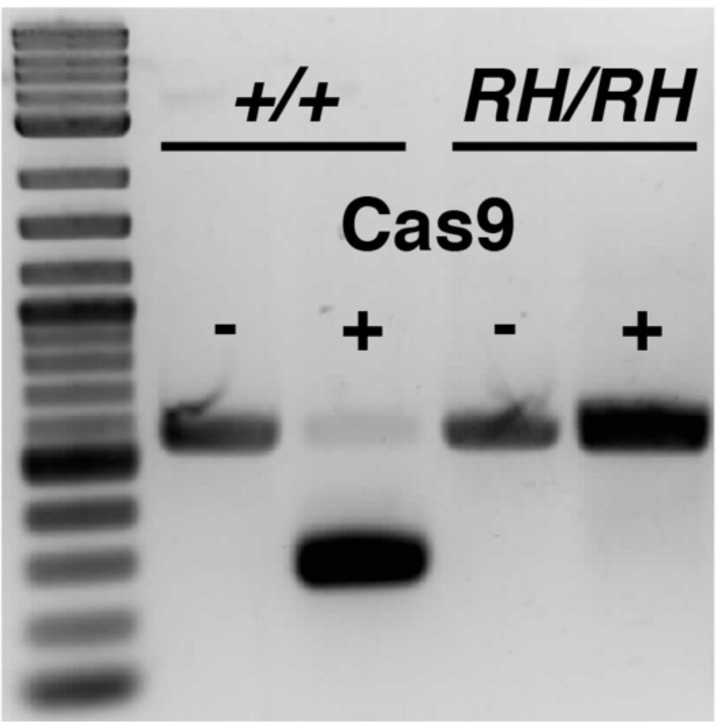
*Chd2* crRNA specifically recognizes wildtype allele. PCR was performed using the Chd2-RH_F/R primer pair with DNA purified from either wildtype or *Chd2^R1684H/R1684H^* mice and the PCR product was used as a template for *in vitro* digestion assays with or without Cas9.

**Extended Data Figure 4.**
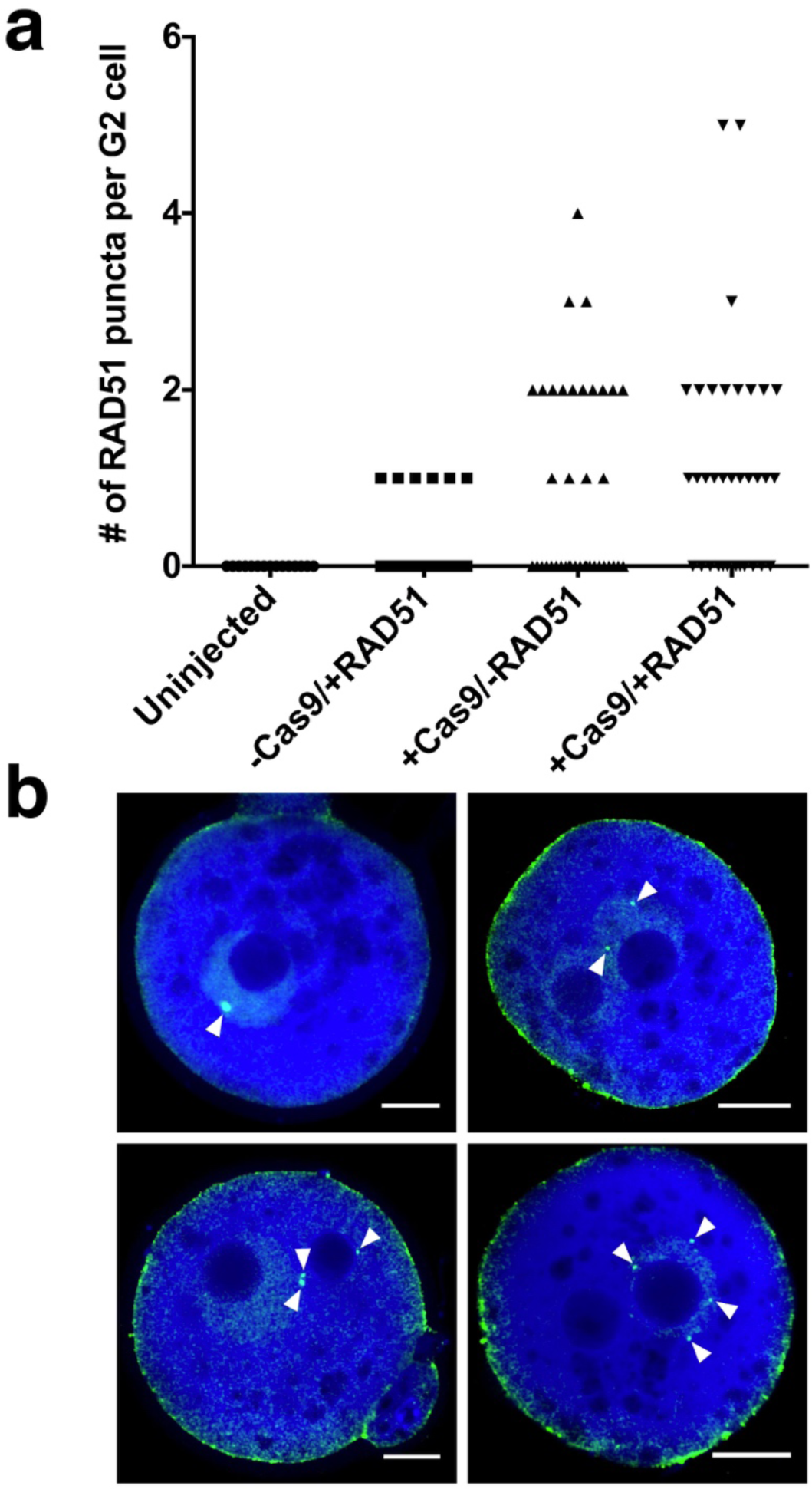
Cas9 injection alters the number of RAD51 puncta in G2. **a,** Quantification of RAD51 puncta number observed in control G2 zygotes and cells injected with RAD51 alone, Cas9/crRNA/tracrRNA alone, or Cas9/crRNA/tracrRNA and RAD51. Each point represents a single cell. **b,** Representative images of cells with one (top left), two (top right), three (bottom left), or four (bottom right) RAD51 puncta (arrowheads indicate puncta; DAPI, blue; RAD51, green; scale=10µm).

**Extended Data Figure 5.**
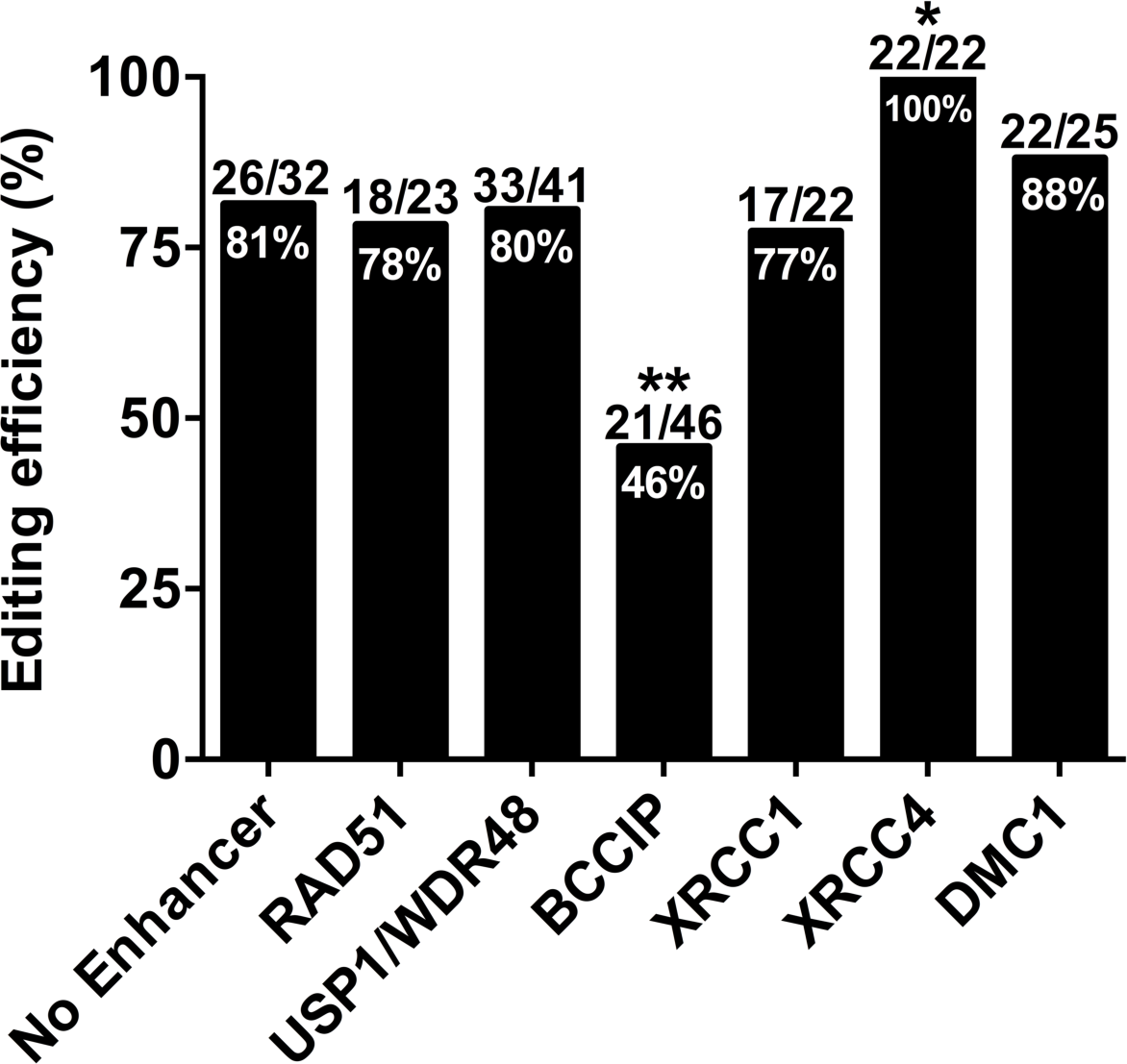
Decreased editing efficiency in BCCIP-injected cells. Editing efficiency observed in IVF-derived embryos injected with Cas9/crRNA/tracrRNA only (no enhancer) or the indicated HR- and DSB repair-associated proteins (BCCIP: p=0.0016, two-tailed chi-square test; XRCC4: p=0.031, two-tailed chi-square test).

**Extended Data Figure 6.**
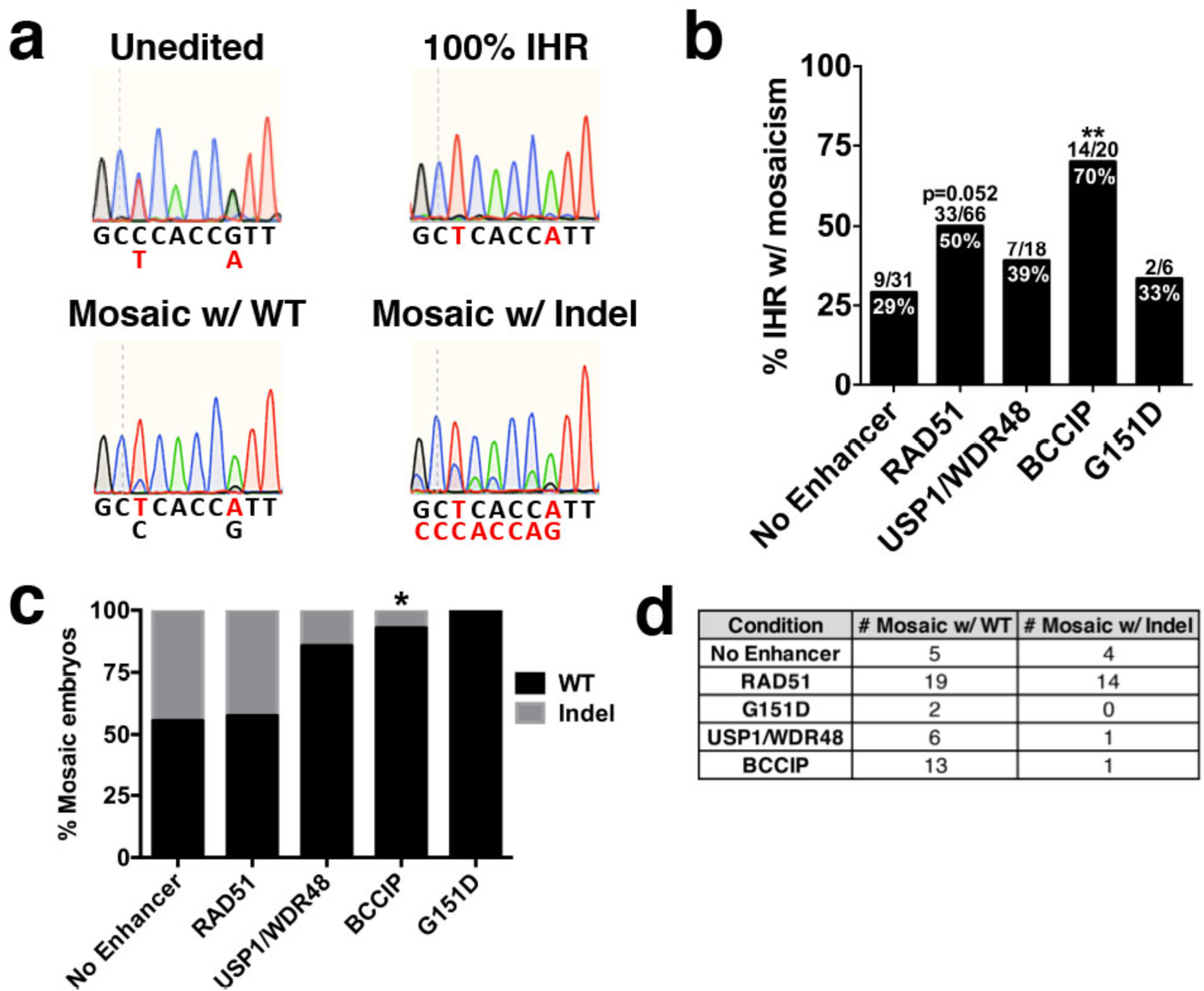
Effects of IHR enhancers on mosaicism. **a,** Representative chromatograms showing examples traces for genotypes characterized as unedited (top left), 100% IHR (top right), mosaic with wildtype (bottom left), and mosaic with indel (bottom right). **b,** Quantification of mosaicism in embryos displaying IHR for all conditions analyzed. Data includes all IVF-derived embryos **(Figs. 3c**, **4c**, and **4d**; BCCIP: p=0.0041, two-tailed chi-square test). **c,** Distribution of types of mosaicism observed in mosaic embryos described in panel b (BCCIP: p=0.034, two-tailed chi-square test). **d,** Numbers for observed mosaicism used to generate panel **c**.

**Extended Data Table 1.** Genotyping Results for Individual F_0_ Pups from CPI and PNI Experiments in Figure 1. Genotyping was performed using the C2RH-Long_F1/R1 primer pair described in Supplementary Table 3.

| Sex | Injection<br>Method | Genotype | Notes |
| --- | --- | --- | --- |
| Female | CPI | RH/RH |  |
| Female | CPI | RH/Indel |  |
| Female | CPI | RH/RH |  |
| Male | CPI | RH/RH |  |
| Male | CPI | RH/RH |  |
| Female | CPI | Indel/Indel | Same Indel |
| Female | CPI | RH/Indel |  |
| Female | CPI | RH/RH |  |
| Male | CPI | Indel/Indel | Different<br>Indels |
| Female | PNI | RH/RH |  |
| Male | PNI | RH/RH |  |
| Male | PNI | RH/RH |  |
| Female | PNI | Indel/Indel | Different<br>Indels |
| Male | PNI | Indel/Indel | Same Indel |
| Male | PNI | RH/RH |  |
| Male | PNI | Indel/Indel | Same Indel |
| Female | PNI | Indel/Indel | Same Indel |
| Female | PNI | RH/RH |  |

**Extended Data Table 2.** Genotype-phenotype analyses of Tyr^C89S^ pups from Figure 2b-e. Phenotype analysis (coat color) and corresponding genotyping data from genomic DNA purified from albino tissue (if available).

| Condition | Coat Color | Genotype |
| --- | --- | --- |
| -RAD51 | White | C89S/C89S |
| -RAD51 | White | C89S/Indel |
| -RAD51 | White | C89S/Indel |
| -RAD51 | White | C89S/Indel |
| -RAD51 | White | C89S/Indel |
| -RAD51 | White | C89S/Indel |
| -RAD51 | White | C89S/Indel |
| -RAD51 | White | Indel/Indel (Different) |
| -RAD51 | White | Indel/Indel (Different) |
| -RAD51 | White | Indel/Indel (Different) |
| -RAD51 | White | Indel/Indel (Different) |
| -RAD51 | Mixed | C89S/C89S |
| -RAD51 | Mixed | C89S/C89S |
| -RAD51 | Mixed | C89S/Indel |
| -RAD51 | Mixed | C89S/Indel |
| -RAD51 | Mixed | C89S/Indel |
| -RAD51 | Mixed | C89S/Indel |
| -RAD51 | Mixed | C89S/Indel |
| -RAD51 | Mixed | Indel/Indel (Different) |
| -RAD51 | Mixed | Indel/Indel (Different) |
| -RAD51 | Mixed | Indel/Indel (Different) |
| -RAD51 | Black | C89S/+ |
| -RAD51 | Black | C89S/+ |
| -RAD51 | Black | Indel/+ |
| -RAD51 | Black | Indel/+ |
| +RAD51 | White | C89S/C89S |
| +RAD51 | White | C89S/C89S |
| +RAD51 | White | C89S/C89S |
| +RAD51 | White | C89S/C89S |
| +RAD51 | White | C89S/C89S |
| +RAD51 | White | C89S/C89S |
| +RAD51 | White | C89S/Indel |
| +RAD51 | White | C89S/Indel |
| +RAD51 | White | C89S/Indel |
| +RAD51 | White | Indel/Indel (Same) |
| +RAD51 | Mixed | C89S/C89S |
| +RAD51 | Mixed | C89S/C89S |
| +RAD51 | Mixed | C89S/C89S |
| +RAD51 | Mixed | C89S/C89S |
| +RAD51 | Mixed | C89S/C89S |
| +RAD51 | Mixed | C89S/C89S |
| +RAD51 | Mixed | C89S/Indel |
| +RAD51 | Mixed | C89S/Indel |
| +RAD51 | Mixed | C89S/Indel |
| +RAD51 | Mixed | Indel/Indel (Different) |
| +RAD51 | Mixed | Indel/Indel (Same) |
| +RAD51 | Black | C89S/+ |
| +RAD51 | Black | C89S/+ |
| +RAD51 | Black | C89S/+ |
| +RAD51 | Black | Indel/+ |
| +RAD51 | Black | Indel/+ |
| +RAD51 | Black | Indel/+ |

**Extended Data Table 3.** Chi-square analyses of immunocytochemistry data from Figure 4b.

| Condition | Positive | Negative | Total | p-Value vs. Uninjected | p-Value vs. RAD51-only | p-Value vs. Cas9-only | p-Value vs. Cas9 + RAD51 |
| --- | --- | --- | --- | --- | --- | --- | --- |
| Uninjected | 0 | 15 | 15 | N/A | 0.0677 | 0.002 | <0.0001 |
| RAD51-only | 6 | 25 | 31 | 0.0677 | N/A | 0.016 | 0.0003 |
| Cas9-only | 17 | 22 | 39 | 0.002 | 0.016 | N/A | 0.0646 |
| Cas9 + RAD51 | 22 | 14 | 36 | <0.0001 | 0.0003 | 0.0646 | N/A |

